# Characterization and significance of extracellular polymeric substances, reactive oxygen species, and extracellular electron transfer in methanogenic biocathode

**DOI:** 10.1101/2020.11.12.380824

**Authors:** Basem S. Zakaria, Bipro Ranjan Dhar

**Author notes:** Corresponding author (B.R. Dhar).

## Abstract

The microbial electrolysis cell assisted anaerobic digestion (MEC-AD) holds great promises over conventional anaerobic digestion. This article reports an experimental investigation of extracellular polymeric substances (EPS), reactive oxygen species (ROS), and the expression of genes associated with extracellular electron transfer (EET) in methanogenic biocathodes. The MEC-AD systems were examined using two cathode materials: carbon fibers and stainless-steel mesh. A higher abundance of hydrogenotrophic *Methanobacterium* sp. and homoacetogenic *Acetobacterium* sp. appeared to play a major role in superior methanogenesis from stainless steel biocathode than carbon fibers. Moreover, the higher secretion of EPS accompanied by the lower ROS level in stainless steel biocathode indicated that higher EPS perhaps protected cells from harsh metabolic conditions (possibly unfavorable local pH) induced by faster catalysis of hydrogen evolution reaction. In contrast, EET-associated gene expression patterns were comparable in both biocathodes. Thus, these results indicated hydrogenotrophic methanogenesis is the key mechanism, while cathodic EET has a trivial role in distinguishing performances between two cathode electrodes. These results provide new insights into the efficient methanogenic biocathode development.

## 1. Introduction

The concept of electro-methanogenesis by combining the microbial electrolysis cell (MEC) and anaerobic digestion (AD) has become a promising method for process intensification and improving the stability of digesters ^1–5^. The integrated process is called microbial electrolysis cell assisted anaerobic digester (MEC-AD). In MEC-AD systems, methane can be produced via multiple pathways, such as (1) direct electron transfer from the cathode to electrotrophic methanogens coupled with CO_2_ reduction to methane, and (2) hydrogenotrophic methanogenesis of H_2_ produced via cathodic hydrogen evolution reaction (HER) ^4–7^. Moreover, methane can also be produced via direct interspecies electron transfer (DIET) between electroactive bacteria (EAB) and electrotrophic methanogens in cathode and anode electrodes ^7–9^. Nonetheless, a considerable portion of methane would still be generated via conventional acetoclastic and hydrogenotrophic methanogenesis pathways.

The activity of anodic EAB was identified as one of the key factors for boosting the methanogenesis process in MEC-AD systems. EAB can outcompete acetoclastic methanogens due to faster growth kinetics ^10^, and divert electrons from acetate to anode via extracellular electron transport (EET). The transferred electrons can be utilized for hydrogen production via a cathodic HER. Thus, fast-growing hydrogenotrophic archaea can be augmented on the biocathode. Several studies reported enrichment of known hydrogenotrophic methanogens, such as *Methanobacterium* and *Methanobrevibacter* in the biocathode ^8–11^. Thus, MEC-AD can provide faster methanogenesis rates compared to conventional anaerobic digesters. Furthermore, MEC-AD systems could provide better process stability due to the faster utilization of volatile fatty acids (VFAs) by EAB ^10,12^. The accumulation of VFAs has been widely reported as a critical factor influencing failure or process instability of digesters operated at high organic loading rates ^13–15^. A few studies also suggested that MEC-AD systems could provide better resilience to inhibitory compounds (e.g., phenol, ammonia, etc.) and decline of digester performance at lower temperatures ^16,17^. Thus, MEC-AD systems can provide numerous benefits over conventional AD.

Despite significant research efforts towards developing MEC-AD systems, studies exploring the significance of extracellular polymeric substance (EPS) in biocathode are limited. Biofilm EPS can have many functions, including attachment of cells to solid surfaces, maturation of biofilm structures, and protection of cells from harsh environmental conditions ^18–20^. A few recent studies validated the significance of EPS in EET within electroactive anode biofilms ^20–23^. In general, EPSs are composed of proteins, extracellular DNA (eDNA), humic acids, polysaccharides, etc., that are secreted by microbes in pure and mixed cultures ^19,20^. Notably, humic acids, eDNA, and heme-binding proteins showed redox properties, serving as immobilized electron carriers in electroactive biofilms ^20,22,23^. Interestingly, EPS extracted from anaerobic digesters also exhibited redox properties and identified as an essential route for DIET between syntrophic bacteria and methanogens ^24,25^. As direct electron transport from the cathode-to-methanogen and bacteria-to-methanogens can promote electro-methanogenesis in the biocathode, it can be assumed that biocathode EPS can potentially be linked with MEC-AD performance. However, to the best of the authors’ knowledge, reports on biocathode EPS characteristics and expressions of EET genes in MEC-AD systems are still scarce.

The optimization of applied voltage/potential and inoculation method has been broadly investigated to enrich a syntrophically balanced microbiome for MEC-AD systems ^2,26^. Previous studies also substantiated the importance of persuasive system design ^27–29^. Particularly, cathode materials with low overpotential, large surface area, and good conductive properties were found to play a deterministic role in MEC-AD performance ^3,30–32^. Carbon-based electrodes, such as carbon fiber, carbon cloth, and carbon brush, have been mostly employed in previous studies due to their high surface area and biocompatibility properties ^3,30,32^. Furthermore, low-cost 3D porous carbon-based composite materials have been developed for the efficient growth of biofilms ^33–35^. However, carbon-based electrodes provide slow catalysis for cathodic HER, which seems to be critical for enriching hydrogenotrophic methanogens ^1,30,36^. Some previous studies employed metal catalysts (e.g., nickel, platinum, etc.) on carbon electrodes to accelerate HER ^1,30,36^, while these catalysts are still expensive. In contrast, non-precious metal electrodes, such as stainless steel, have shown an excellent low-cost alternative ^11,31,37–39^. However, to date, limited information is available on how carbon and metal-based electrodes shape biocathode structures in terms of EPS, expression of EET genes, and microbial communities.

Considering the research gaps mentioned above, the main goal of this study was to provide fundamental insights into the EPS characteristics and EET genes in methanogenic biocathode. The novelty of this study is two folds. First, this study presents, for the first time, a comprehensive characterization and significance of EPS and expression of EET genes for methanogenic biocathode. Second, underlying mechanisms of methanogenesis performance with carbon and metal cathodes were evaluated with a multifaceted approach combining molecular biology, microscopic and electrochemical tools.

## 2. Materials and methods

### 2.1. Experiment

Two single chamber MEC-AD systems (working volume of 360 mL), constructed with plexiglass, were used in this experiment. Carbon fibers (2293-A, 24A, Fibre Glast Development Corp., Ohio, USA) fixed onto a stainless-steel frame (which was not exposed to the liquid medium) was used as an anode electrode in both reactors. A similar carbon fiber module was used as a cathode electrode in one reactor (referred to as ‘CF-CF’), and a stainless-steel mesh (304 stainless steel, McMASTER-CARR, USA) was used as a cathode in the other reactor (referred to as ‘CF-SS’). The specific surface area provided by the stainless-steel electrode was 4.2 m^2^/m^3^. Considering every single filament in the carbon fiber bundle (Fig. S1), the specific surface area of the carbon fibers module was estimated at 3609 m^2^/m^3^. Moreover, considering all filaments in a bundle as a single fiber, the surface area was estimated at 41 m^2^/m^3^. The detailed calculation of the specific surface areas is provided in the Supporting Information. The Ag/AgCl reference electrode (MF-2052, Bioanalytical System Inc., USA) was positioned close (<1 cm) to the anode electrode.

Both reactors were inoculated with mesophilic anaerobic digester sludge and effluent from a dual-chamber mother MEC operated with sodium acetate as an electron donor for >12 months. Initially, sodium acetate (1600±55 mg COD/L) supplemented phosphate buffer (50 mM) and trace minerals were used as a substrate. The details of the trace minerals can be found elsewhere ^2^. Both reactors were operated with acetate in fed-batch mode until peak current densities reached ∼77 A/m^3^. Then, the substrate was switched to glucose (2150±31 mg COD/L), while buffer and trace minerals composition remained the same. With glucose, reactors were operated for about six months under batch mode in repetitive cycles. The biogas produced from the reactors were collected in gasbags. A decrease in daily methane production to <3 mL was considered an indicator for replacing the substrate medium. During experiments, the anode potential was fixed at −0.4 vs. Ag/AgCl with a potentiostat (Squidstat Prime, Admiral Instruments, Arizona, USA). This anode potential was selected to enrich and maintain kinetically efficient EAB as suggested in the literature ^40–42^. The reactors were operated at room temperature (21±1°C) with continuous mixing (130±5 rpm) of the liquid medium with magnetic stirrers.

### 2.2. EPS and ROS analyses

For EPS analysis, biomass samples were washed three times with 0.1 M PBS (pH 7.4), and then the supernatant was removed after centrifugation at 3000 × g for 15 min at 21°C. EPS extraction was performed using two methods: cation-exchange resin (CER) and heating method. Applying cation-exchange resins and heating methods was highly efficient in several previous studies for the EPS extraction from biofilms, particularly carbohydrates, proteins, and eDNA ^20,23,43–45^. In addition, the pellets were collected to examine the cell lysis interference using a Glucose-6-Phosphate Dehydrogenase kit (Sigma-Aldrich, USA). The details of EPS extraction and analytical methods for various EPS components (proteins, carbohydrates, eDNA, heme-binding proteins, and uronic acid) are provided in the Supporting Information. These five major EPS components were selected based on the EPS components previously found in electroactive anode biofilms ^20,46–48^. Furthermore, these EPS components were also found in archaeal biofilms in conventional anaerobic biofilm reactors ^49,50^. Both EPS extraction methods (CER and heating method) provided similar results (Table S1). Hence, we reported the results from the CER method. Confocal laser scanning microscopy (CLSM) was used to visualize and examine the EPS structure (for method, see Supporting Information). The quantitative analysis of EPS biovolumes and intensities was carried out using biofilm image processing COMSTAT software. For electrochemical characteristics, cyclic voltammetry (CV) of extracted EPS from biocathodes was performed. 2 mL of extracted EPS was transferred to an electrochemical cell having screen printed electrodes (A-AD-GG-103-N, Zimmer and Peacock Ltd., Royston, UK). The working electrode potential was ramped between − 0.8 and 0.8 V vs. Ag/AgCl at a scan rate of 1 mV/s using a potentiostat (Squidstat Prime, Admiral Instruments, USA); the current was recorded every 1 second.

The ROS levels in biofilms were visualized using CLSM. We collected different parts of electrodes, then washed them three times using 0.1 M PBS to remove any debris. Samples were stained with 2’,7’-Dichlorofluorescein diacetate (10 µM) (Thermo Fisher, USA) for one hour. The visualization (Fig. S2) was then performed with excitation and emission wavelength of 495 nm and 520 nm, respectively. The quantification of ROS levels was then performed using image processing COMSTAT software. Microscopic visualization of biofilms was performed using a scanning electron microscope (SEM, Carl Zeiss, Cambridge, UK). Several images from different locations of electrodes were captured. The detailed protocol could be found elsewhere ^19^.

### 2.3. Microbial communities and gene expression analyses

For microbial analyses, genomic DNA was extracted from the anode and cathode biofilms using PowerSoil^®^ DNA Isolation Kit (MoBio Laboratories, Carlsbad, USA) according to the manufacturer’s instructions. The purity and concentration of DNA were measured with Nanodrop spectrophotometer (Model 2000C, Thermo Scientific, USA). The extracted DNA was stored immediately at −70 °C prior to the sequencing. Illumina Miseq Sequencing was performed by the Research and Testing Laboratory (Lubbock, TX, USA) targeting 16S rRNA gene using bacterial primers 341F: 5’ CCTACGGGNGGCWGCAG 3’ and 805R: 5’ GACTACHVGGGTATCTAATCC 3’ ^51,52^, archaeal primers 517F: 5’ GCYTAAAGSRNCCGTAGC 3’ and 909R: 5’ TTTCAGYCTTGCGRCCGTAC 3’ and specific *mcr*A archaeal primers *mcr*Af: 5’ GGTGGTGTMGGATTCACACARTAYGCWACAGC 3’ and *mcr*Ar: 5’ TTCATTGCRTAGTTWGGRTAGTT 3’ ^53^.

For evaluating microbial diversity, the nucleotide sequence reads were sorted out using a data analysis pipeline. Short sequences, noisy reads and chimeric sequences were removed through a denoising step and chimera detection, respectively. Then, each sample was run through the analysis pipeline to establish the taxonomic information for each constituent read. Microbial taxonomy was assigned using the Quantitative Insights Into Microbial Ecology (QIIME) pipeline (http://qiime.org ^54^). Principal component analysis (PCA) of cathodic microbial communities was conducted using weighted Unifrac metrics to show the relation between genera and PCs. The expressions of EET genes (i.e., *pil*A, *omc*B, *omc*C, *omc*E, *omc*Z, and *omc*S) were also quantified (for details and method, see Supporting Information). The primers and design methods are listed in Table S2.

### 2.4. Analytical methods and statistical analysis

Current and applied voltage/potential were recorded every 4.8 min using a computer connected with the potentiostat. The chemical oxygen demand (COD) was measured with HACH method using UV-spectrophotometer (Model DR 3900, HACH, Germany). The volatile fatty acids, VFAs (acetate, propionate, and butyrate) concentrations were measured with an ion chromatography (DionexTM ICS-2100, Thermo Scientific, USA) ^42^. Electrochemical impedance spectroscopy (EIS) was performed for both reactors using a multi-channel VSP potentiostat (VSP, Bio-Logic Science Instruments, France). The detailed methodology is provided in the Supporting Information. The biogas produced from reactors was collected with 500 mL gas bags. The composition of biogas (i.e., methane content) was analyzed with a gas chromatograph (7890B, Agilent Technologies, Santa Clara, USA) equipped with a thermal conductivity detector and two columns (Molsieve 5A and Hayesep). To reveal the statistical difference between the results collected from two reactors, the student’s paired t-test (JMP^®^ Software, SAS, US) was used.

## 3. Results and discussion

### 3.1. MEC-AD performance

The performance of the two configurations was compared based on volumetric current density and methane productivity. As shown in Fig. S3, the maximum current density from the CF-SS reactor reached 34.1±0.3 A/m^3^, which was significantly higher (p=0.01) than CF-CF (27.6±0.2 A/m^3^). Although the methane generation patterns were comparable in both reactors (Fig. 1), CF-SS showed higher (p=0.03) daily methane production than CF-CF throughout the batch cycle. The total cumulative methane production was substantially higher in CF-SS (179.5±6.7 vs. 100.3±7.9 mL CH_4_; p=0.01). Both reactors used carbon fiber as the anode electrode and were operated under identical operating conditions (e.g., mixing speed, substrate, inoculation, etc.). Hence, the differences in system performance could be closely tied to the difference in the cathode electrode. As discussed later, stainless steel mesh cathode in CF-SS facilitated denser biofilms formation with more methanogenic biomass.

**Fig. 1.**
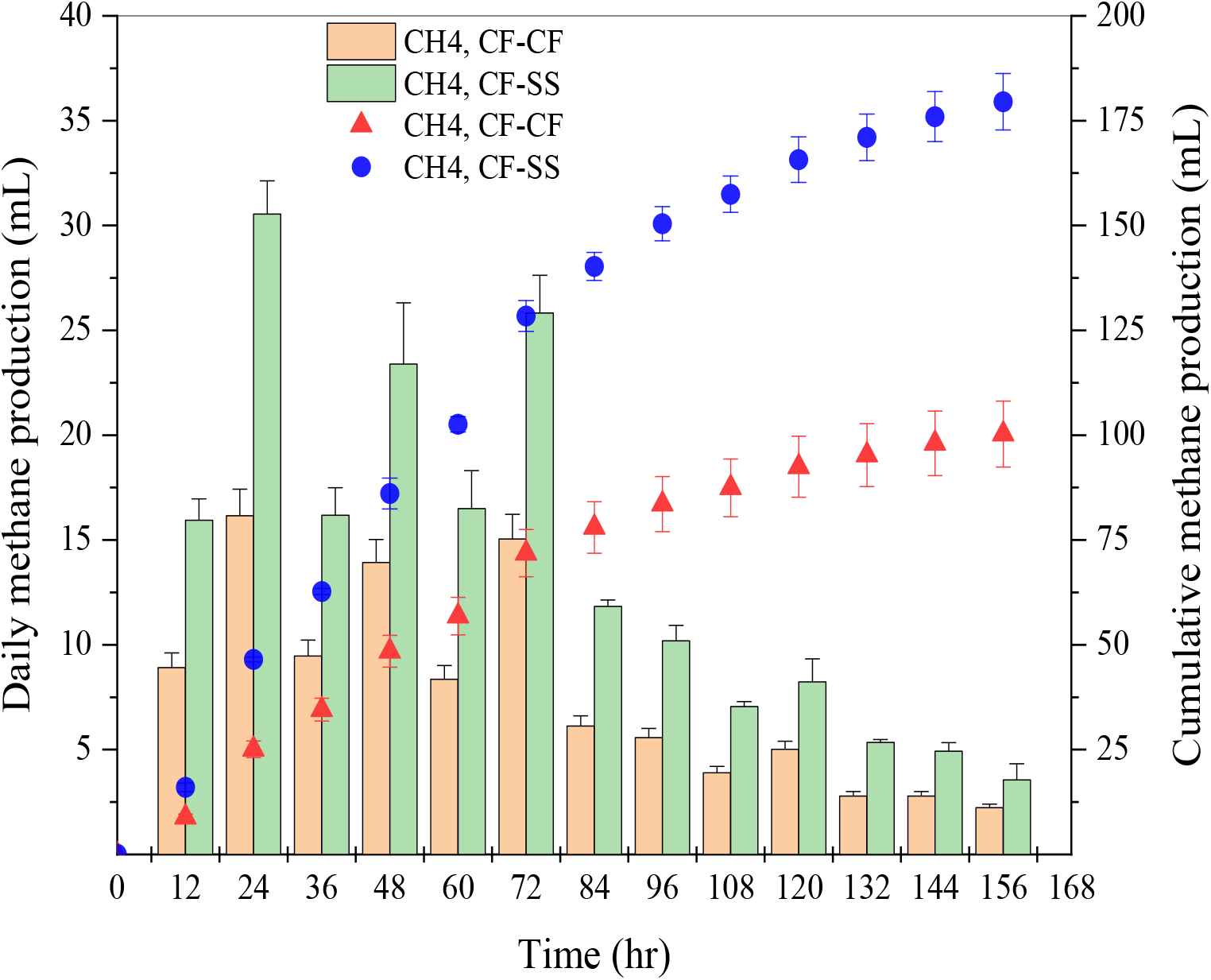
Methane production from CF-CF and CF-SS reactors. The error bars indicate the standard deviation of three replicates (n = 3).

Anode electrodes providing high specific surface areas have been efficient for enhancing the performance of various bio-electrochemical systems ^3,28,30–32^. Therefore, carbon-based electrodes, such as carbon brush, activated carbon, have also been widely used for various biocathode applications, including electro-methanogenesis ^3,30,32^. Notably, the rough surface of carbon fiber was found to be efficient for developing EAB biofilms ^55^. Although we cannot rule out that different textures (diameter of carbon fiber and stainless steel wire) could also lead to distinct colonization of biomass ^56,57^, the specific surface area provided by electrodes is often considered a critical factor. This study shows that stainless steel cathode having a relatively lower specific surface area than carbon fibers (4.23 vs. 3609 m^2^/m^3^) resulted in a superior methanogenic activity. It has been previously suggested that the agglomeration of fibers in the liquid phase could reduce the available specific surface area for biofilms formation ^58^. Nonetheless, considering all carbon fiber filaments in a bundle as a single fiber, the specific surface area provided by the carbon fiber was still higher than stainless-steel (4.23 vs. 41 m^2^/m^3^). In general, carbon-based electrodes are considered inferior catalysts for HER than metal and carbon-metal composite electrodes ^11,31,37– 39^. Previous MEC-AD studies substantiated the role of hydrogenotrophic methanogenesis. It is also reasonable that acetoclastic methanogens would likely be washed out at low residence time (<7 days) used in this study ^7,59^. EIS analysis also indicated that stainless-steel biocathode could reduce various intrinsic internal resistances in CF-SS compared to CF-CF (see Supporting Information). As shown in the Nyquist plot (see Fig. S4), the overall internal impedance of CF-SS (52.62 Ω) was lower than that of CF-CF (77.28 Ω). Thus, stainless-steel cathode largely influenced the internal resistances, which influenced the HER kinetics and, ultimately, growth and activities of hydrogenotrophic methanogens. Previous studies also suggested that lower ohmic resistance in MECs could provide faster HER kinetics ^60,61^. Thus, the inferior methane recovery from the CF-CF reactor than the CF-SS reactor was likely due to the inferior HER on carbon fibers and subsequent hydrogenotrophic methanogenesis.

### 3.2. Organics removal and VFAs profiles

The effluent COD concentration from CF-SS (215±2.8 mg/L) was considerably lower (p=0.001) than that of CF-CF (382±3.0 mg/L) (Fig. S5a). Correspondingly, COD removal efficiency in CF-SS (89.9±0.5%) was significantly higher (p=0.012) than that of CF-CF (83±1.7%). Fig. S5b and S5c show the VFAs profiles during batch operation. For both reactors, the acetate concentrations were relatively higher than propionate and butyrate throughout the operational period. The CF-SS reactor showed the highest acetate concentration of 439±2 mg COD/L, while propionate (94±0.1 mg COD/L) and butyrate (61±0.3 mg COD/L) concentrations were relatively lower. In contrast, CF-CF exhibited the highest acetate concentration of 320±0.4 mg COD/L, which was lower than that observed in CF-SS. Propionate concentrations were relatively higher in CF-CF, with the highest concentration of 118±0.4 mg COD/L. The highest butyrate concentration (64.8±1.3 mg COD/L) in CF-CF was comparable to CF-SS (61±0.3 mg COD/L). CF-SS also showed a lower accumulation of VFAs in the final effluent than CF-CF (52.6±0.5 vs. 133.5±1.0 mg COD/L; p=0.005).

Throughout the batch operation, propionate concentrations in CF-SS remained relatively lower than those observed in CF-CF, indicating faster conversion of propionate in the CF-SS. The fermentation of propionate to acetate is a vital process towards anodic respiration (by EAB) and acetoclastic methanogenesis. However, propionate fermentation to acetate is energetically unfavorable in terms of Gibbs free energy ^62^. Thus, maintaining lower hydrogen partial pressure would be critical for propionate fermentation to acetate. Even though stainless-steel cathode would be expected to provide superior HER than carbon fibers ^11,38^, no hydrogen was detected in biogas from both reactors. This might be due to the rapid consumption of hydrogen by hydrogenotrophic methanogens, as suggested in previous studies ^11,63^. Moreover, enhanced homoacetogenic activity (H_2_ + CO_2_ → acetate) could assist in maintaining lower hydrogen partial pressure in biocathode ^11,63^. Microbial community analysis also coincided with these notions (discussed later). Thus, the VFA profiles suggest that the microbiome in CF-SS more rapidly utilized hydrogen produced via fermentation and cathodic HER.

### 3.3. EPS characteristics

As shown in Fig. 2a, the EPS composition of anode biofilms in both reactors was quite similar and was not affected by the different cathode materials used. Protein was found as the major EPS component in anode biofilms, consistent with recent reports on EPS composition in pure culture *Geobacter* biofilms ^23,46^. *Geobacter* species were also abundant in anode biofilms in both reactors in this study (discussed later). The concentrations of major EPS components (carbohydrates, proteins, and hemes) in the cathode biofilms in CF-SS were higher than those of CF-CF. Notably, carbohydrates and proteins in cathodic EPS were markedly higher in CF-SS than CF-CF (carbohydrates: 52.2±0.2 vs. 25.8±0.5 mg/cm^2^; proteins: 212.8±3.4 vs. 170±1.3 mg/cm^2^). The heme-binding proteins, uronic acid, and eDNA also showed the same patterns. Overall, cathodic biofilms developed on the stainless-steel electrode exhibited markedly higher EPS levels (p=0.03).

**Fig. 2.**
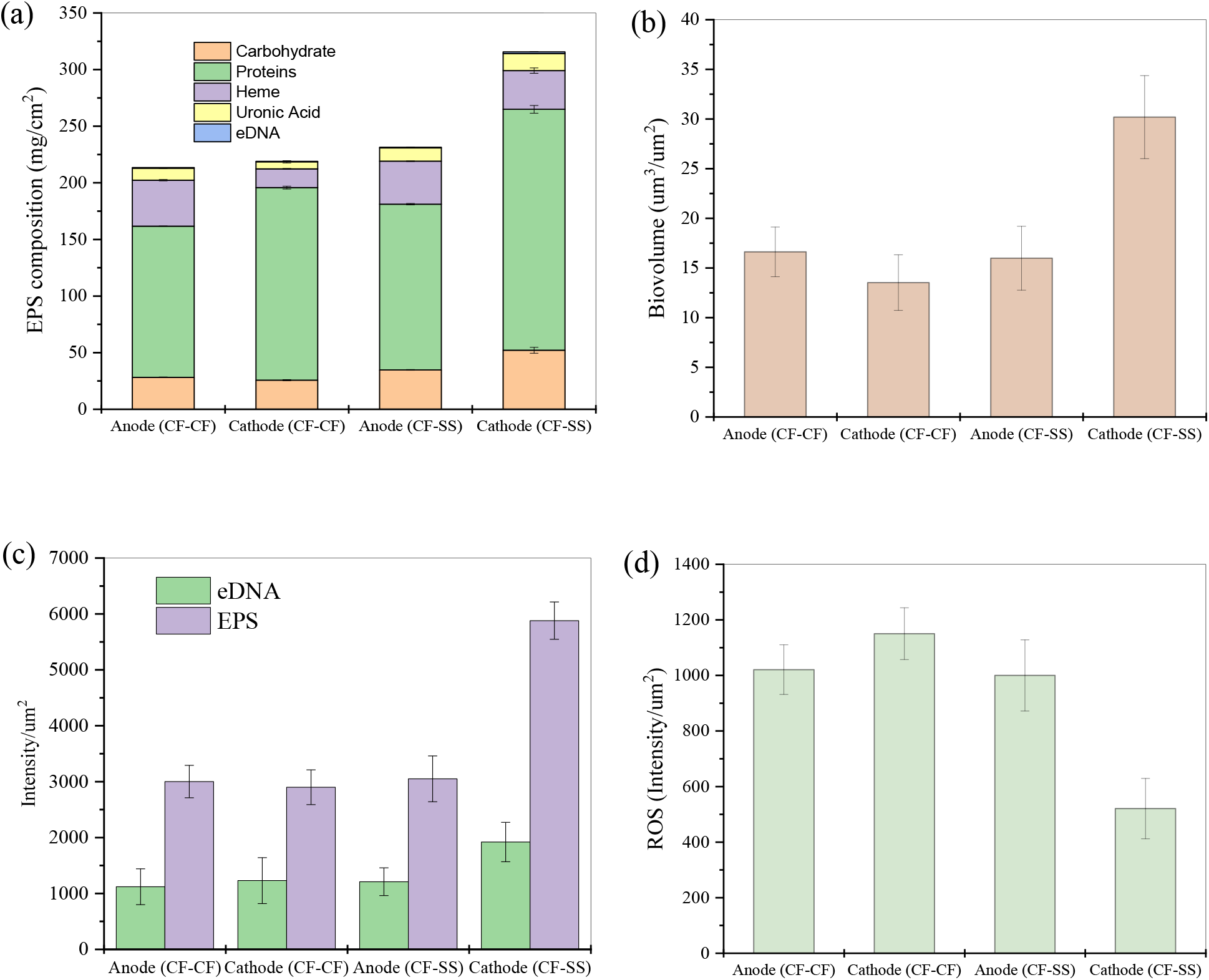
EPS levels in biofilms (a), EPS quantitative analysis using CLSM; biovolume (b), and fluorescence intensity (c), and reactive oxygen species (ROS) intensities (d) of CF-CF and CF-SS reactors. Note. The error bars indicate the standard deviation of three replicates (n = 3).

Moreover, electrode surfaces were visualized with CLSM (Fig. 3). The CLSM images showed that EPS was more uniformly distributed on the stainless-steel biocathode in CF-SS than the carbon fiber electrodes in both reactors. The biovolume of cathode biofilms in CF-SS was estimated at 30.2±4.2 µm^3^/µm^2^, which was two times higher than that estimated for cathode biofilms in CF-CF (13.5±2.8 µm^3^/µm^2^) (Fig. 2b). The biovolumes estimated for anode biofilms in both reactors were comparable (p=0.007). The intensities of EPS and eDNA were also quantified (Fig. 2c). Like estimated biovolume, EPS and eDNA intensities in stainless steel biocathode were higher than those estimated for carbon fiber biocathode (p=0.008). Simultaneously, EPS and eDNA intensities were comparable for anode biofilms in both reactors (p=0.20). Thus, CLSM imaging and COMSTAT analysis further confirmed that the stainless-steel biocathode resulted in the highest EPS production. The SEM imaging of biofilms also corroborated these results (Fig. S6). The biofilms did not fully cover the surfaces of carbon fibers, while biofilms grown on stainless steel cathode in CF-SS were evenly denser than anode/cathode biofilms grown on carbon fiber electrodes. The anode/cathode biofilms grown on carbon fibers exhibited substantial heterogeneity. In contrast, a large secretion of EPS could accelerate the surface attachment of cells on the stainless-steel.

**Fig. 3.**
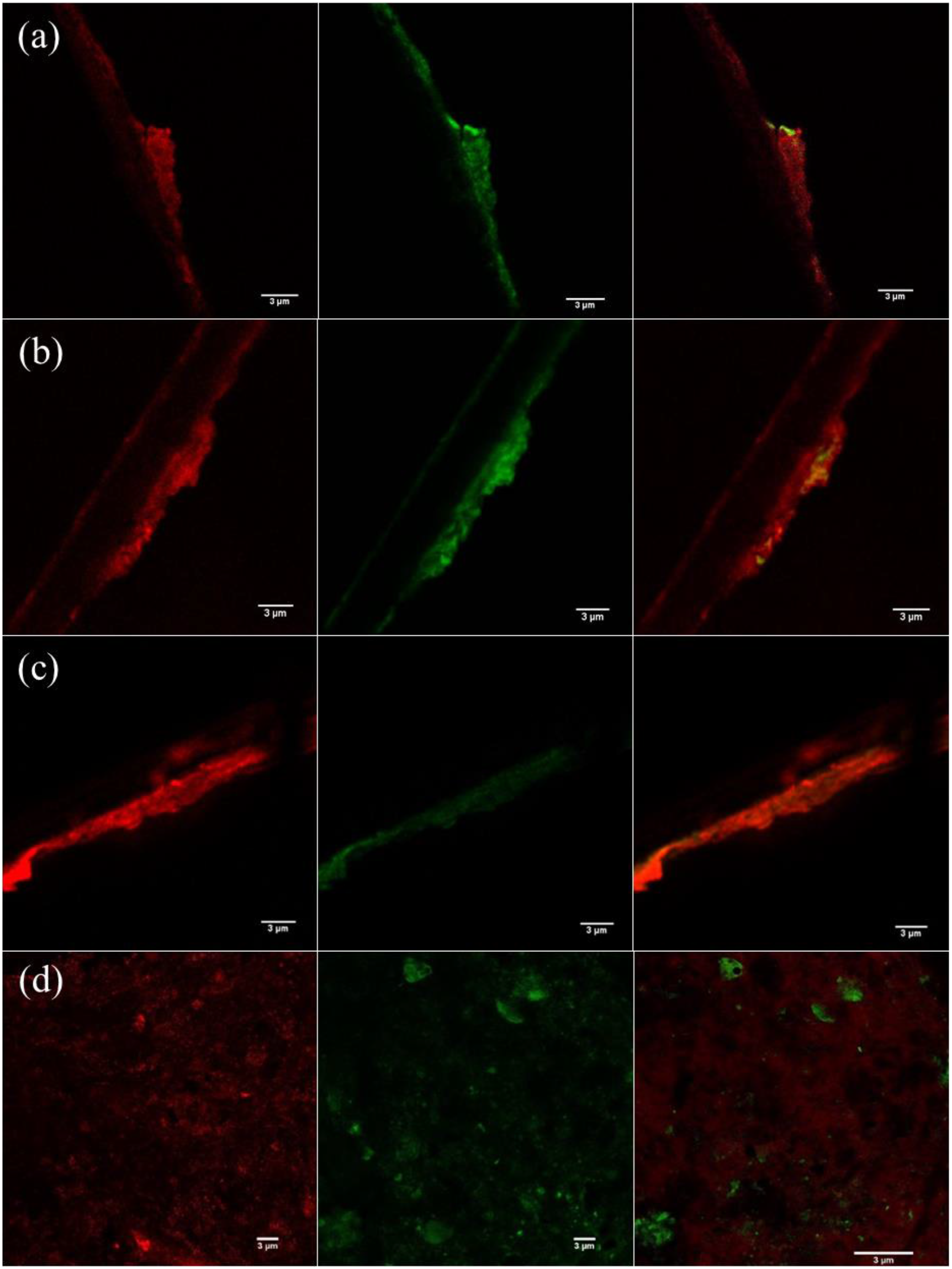
Representative confocal microscopic images of EPS with 3 µm scale; anode (CF-CF) (a), cathode (CF-CF) (b), anode (CF-SS) (c), and cathode (CF-SS) (d). The green color represents eDNA and the red color indicates EPS.

Studies on the EPS in electroactive biofilms received less attention and primarily focused on understanding their role in anodic EET. A few reports revealed redox-active features of anodic EPS in model EAB biofilms (e.g., *Geobacter sulfurreducens, Shewanella oneidensis*, and *Pseudomonas putida*) ^20,23,46^. Notably, higher levels of proteins in anode biofilms were correlated with higher EET efficiency. In this study, despite differences in volumetric current densities, both EPS composition and concentrations were quite similar in anode biofilms in both reactors. Instead, the difference in cathodic EPS levels was likely linked to current densities and methane productivity. As mentioned earlier, EPS can serve as immobilized redox cofactors (i.e., electron carriers) for facilitating EET in anodic EAB biofilms ^20,23^. EAB can also regulate EPS generation to balance EET and protect cells ^48^. The existing literature provides limited information on the roles of EPS in methanogenic biocathode. However, a few reports suggested that EPS could play similar roles (EET and cell protection) in archaeal biofilms in conventional digesters in the presence of conductive additives ^24,64^. Interestingly, a recent study demonstrated that the addition of iron-based conductive materials in conventional anaerobic bioreactors could enhance redox-active EPS contents in methanogenic biomass ^24^, which was positively correlated with methanogenesis rates. Conductive materials promote the syntrophic DIET from bacteria to archaea and thereby enhance methanogenesis ^65^. Therefore, the CV of biocathode EPS from two reactors was performed to identify their redox activity (Fig. S7) qualitatively.

As shown in Fig. S7, the voltammograms of cathodic EPS extracted from both reactors showed distinct redox peaks, indicating their redox capability. However, redox peaks were observed at different potentials, suggesting that redox properties would be different for EPS extracted from two biocathodes. The peak current from stainless steel biocathode EPS was considerably higher than the EPS extracted from carbon fiber biocathode. This difference could be associated with higher levels of redox-active EPS in stainless steel biocathode, as previously suggested in the literature for anodic EAB biofilms ^20,46^. Despite higher EPS levels in stainless-steel biocathode and differences observed in CV patterns, the expressions of genes associated with EET were comparable in both biocathodes (discussed later). Thus, it can be inferred that redox activities of EPS did not play a decisive role in differentiating between the performances observed from the two systems. Instead, EPS variations might be more associated with the protection of cells from harsh metabolic environments. Nonetheless, future investigation is warranted to reach a more thorough understanding and quantitative characterization of redox properties of EPS.

A recent study reported that the current from anode biofilms was positively associated with EPS protein content and negatively correlated to carbohydrates in EPS ^48^. In this study, both carbohydrates and proteins in EPS were considerably higher in stainless steel biocathode than that of carbon fiber biocathode (see Fig. 2). The secretion of carbohydrates could be associated with harsh environmental conditions ^19,25^ to provide a protective layer and maintain the redox activity of proteins involved in EET ^22,25^. It is possible that enhanced HER in stainless steel cathode could create highly alkaline conditions near the cathode ^7,66,67^, which might induce more EPS secretion. Based on a recent report, hydrogenotrophic methanogenesis could be the dominant pathway under alkaline pH ^68^. As discussed later, stainless steel biocathode also showed a higher abundance of hydrogenotrophic methanogens in this study. Thus, it appeared that higher enrichment of hydrogenotrophic methanogens promoted by faster HER kinetics on stainless steel cathode was possibly associated with higher EPS excretion. While further investigation is needed to get more insights into the function of EPS on electro-methanogenesis, these results suggested that different cathode materials could influence EPS secretion and methanogenic activity due to differences in HER kinetics.

### 3.4. ROS levels

The quantitative measurement of ROS demonstrated a significant difference between biofilms grown on stainless steel and carbon fibers (Fig. 2d). The lowest ROS level was observed for cathode biofilms formed on stainless steel, while ROS levels were very similar in anode/cathode biofilms developed on carbon fibers. Recent studies reported ROS accumulation in anaerobic digesters ^21,69^, while ROS is usually thought to be produced during aerobic metabolism. It has been suggested that unfavorable metabolic conditions (e.g., inhibition by toxicants, pH changes) could lead to ROS accumulation in digesters ^21,69^. ROS accumulation may suppress metabolic activities, leading to the deterioration of digester performance. As we used synthetic glucose medium as a substrate, the potential unfavorable metabolic conditions induced by any toxic compounds can be ruled out. Thus, potential local pH changes by HER can be considered as an unfavorable metabolic condition. The HER in both biocathode can lead to alkaline pH due to protons reduction (2H^+^ + 2e^−^ → H_2_), while effects will likely be more intense on stainless steel cathode ^11,31,37–39^. Thus, the lowest ROS level in stainless steel biocathode suggests that higher EPS levels provided some degree of protection to the cathodic microbiome from potential environmental stress (e.g., local alkaline pH due to superior HER). However, potential mechanisms relating to EPS and ROS levels should be further explored.

### 3.5. Microbial quantity and diversity

Fig. 4 shows the quantitative assessment of microbial communities performed with qPCR. The total microbial cell counts (16S) in anode biofilms in CF-SS were slightly higher than that of CF-CF (9×10^8^ vs. 8×10^8^ cells/cm^2^) (Fig. 4a). An almost similar pattern was observed for cathode biofilms; however, the difference was more prominent (1×10^11^ vs. 6×10^8^ cells/cm^2^). The archaeal cell numbers also showed similar patterns, with the highest archaeal cell numbers for the stainless steel biocathode.

**Fig. 4.**
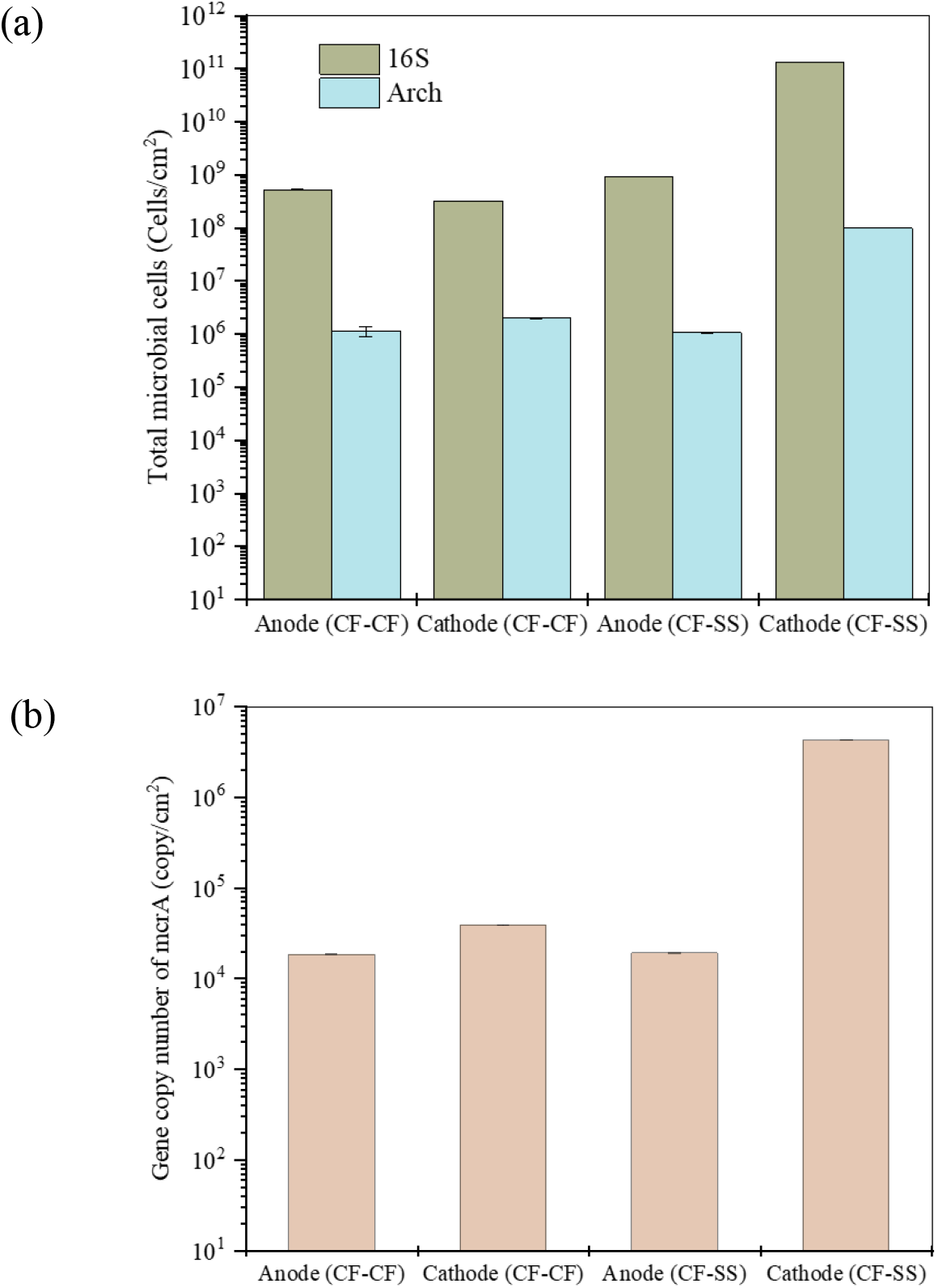
Total cell number using 16s and archaeal primers (a), and *mcr*A gene copies (b). The error bars indicate the standard deviation of three replicates (n = 3).

Furthermore, *mcr*A gene copies were quantified (Fig. 4b), considered a biomarker for hydrogenotrophic methanogenesis ^70^. A few recent reports also confirmed the positive link between *mcr*A gene copies and methanogenesis rates in MEC-AD reactors ^53,70,71^. The highest number of *mcr*A gene copies was observed for the stainless steel biocathode (4×10^6^ cells/cm^2^; 100 times higher than carbon fiber biocathode). The *mcr*A gene copies in anodic biofilms for both systems were comparable. Thus, the higher abundance of *mcr*A gene copies within the stainless steel biocathode corroborated with higher methane productivity in the CF-SS reactor.

The alpha diversity of microbial communities was also estimated (Table S3). The higher values of Chao 1, phylogenetic distance, OTUs, Pielou’s evenness, and Shannon index clearly showed that the richness and diversity indices were relatively higher in CF-SS than CF-CF. Notably, cathode biofilms in CF-SS showed more diversity with the Shannon index of 5.10, as compared to CF-CF (3.95). These results indicated that the stainless-steel electrode persuaded the richness and diversity of the microbial communities.

### 3.6. Microbial community composition, and gene expression

#### 3.6.1. 16S rRNA sequencing

Microbial communities in two reactors were analyzed with specific bacterial, archaeal, and *mcr*A primers. Proteobacteria was the most abundant phylum in anode biofilms in both reactors; however, its relative abundance was much higher in CF-SS (85%) than CF-CF (47%) (Fig. S8). The relative abundances of Bacteroidetes (26%) and Firmicutes (14%) in CF-CF were considerably higher than CF-SS (6% and 4%, respectively). Also, Synergistetes (6%) and Lentisophaerae (4%) were present at slightly higher abundances in CF-CF, while in CF-SS, they were 1% and 3%, respectively. Proteobacteria was also the most abundant in both cathode biofilms; their relative abundances (64%-68%) were also similar. However, the abundance of Bacteroidetes was higher in CF-CF (17%) than CF-SS (6%). On the contrary, the phylum Firmicutes (20%) was the second most abundant in CF-SS, while its abundance in CF-CF was considerably lower (9%).

At the genus level, *Geobacter*, belong to Proteobacteria, was the most abundant in anode biofilms (CF-CF: 22%; CF-SS: 59%) in both systems (Fig. 5a). *Geobacter* is a highly efficient EAB with the capability to facilitate EET from simple organic acids like acetate ^42,48^. In CF-CF, *Bacteroides* was the second most dominant genus (12%), followed by *Enterobacteriaceae* (10%) and *Dysgonomonas* (5%). In contrast, the second abundant genus in CF-SS was *Enterobacteriaceae* (23%), followed by *Dysgonomonas* (3%) and *Victivallis* (3%).

**Fig. 5.**
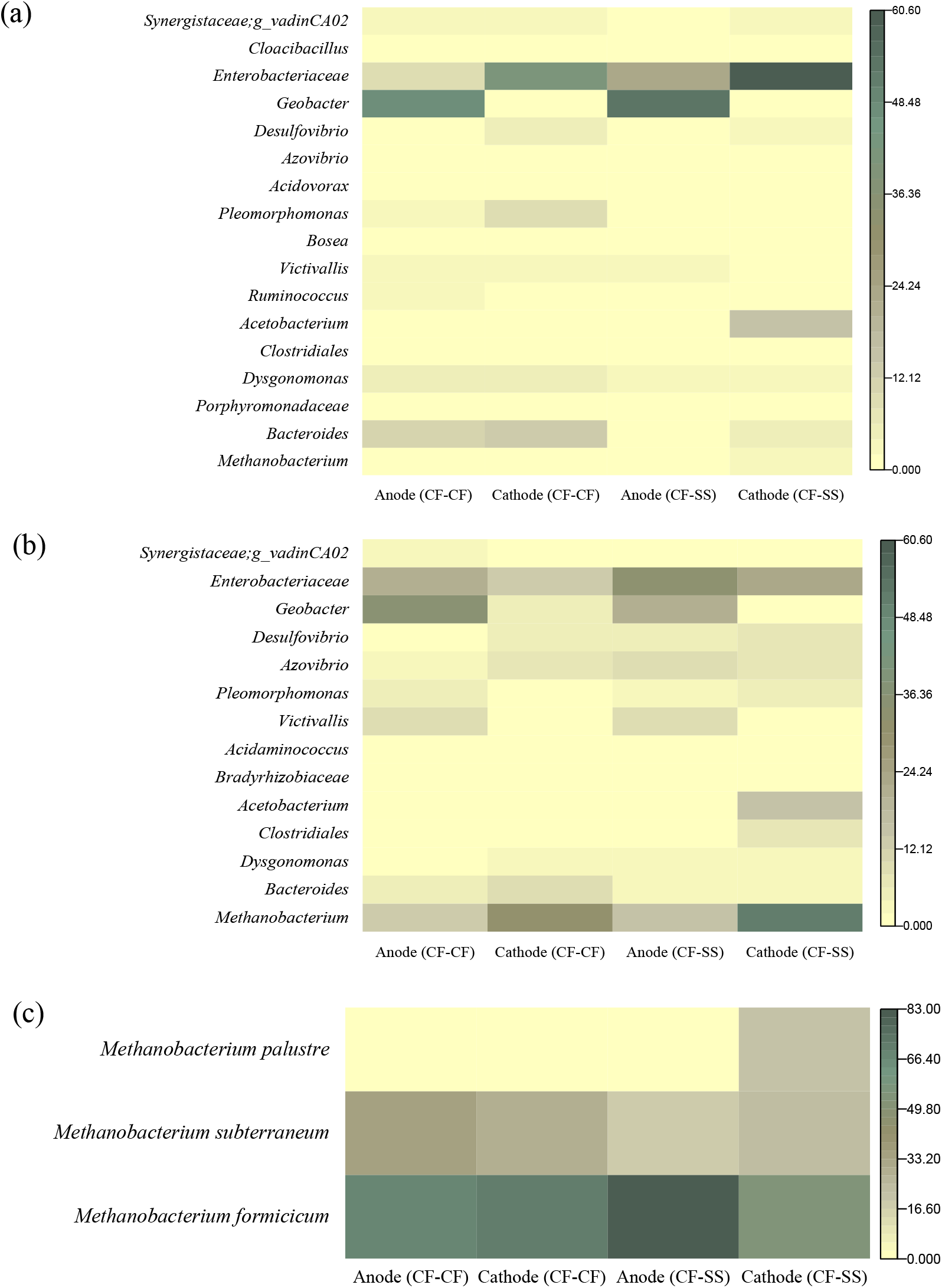
Relative abundance of microbial communities analyzed with bacterial primer (a), archaeal primer (b), and *mcr*A primer (c) at the genus level.

The cathode biofilms in both reactors were dominated by the genus *Enterobacteriaceae* (CF-CF: 42%; CF-SS: 60%). In CF-CF, *Bacteroides* (12%), *Pleomorphomonas* (9%), and *Desulfovibrio* (4%) were the other dominant genera. In contrast, *Acetobacterium* was the second abundant genus (16%), followed by *Bacteroides* (5%), *Dysgonomonas* (3%), and *Desulfovibrio* (2%) in CF-SS. *Acetobacterium*, known homoacetogenic bacteria, can utilize H_2_ and CO_2_ to produce acetate ^31,72^. Then, acetate can be consumed by either acetoclastic archaea or EAB ^7,10,73^. The enrichment of *Acetobacterium* on the stainless-steel biocathode indicates the occurrence of higher catalysis of HER. As mentioned earlier, H_2_ gas has not been observed in the biogas samples. This might be due to the rapid utilization of the generated H_2_ via hydrogenotrophic methanogens and homoacetogenic *Acetobacterium*, as suggested in the literature ^11,63^. The presence of the highest acetate concentration (439±2 mg COD/L) in CF-SS corroborated with a higher abundance of *Acetobacterium*. Moreover, *Acetobacterium* can maintain a lower hydrogen partial pressure to provide thermodynamically favorable conditions for propionate and butyrate fermentation to acetate. This notion is also supported in part by the lower propionate concentrations in CF-SS compared to the CF-CF.

#### 3.6.2. Archaeal and *mcr*A primer sequencing

For the archaeal phylum, relative abundances of Euryarchaeota were 32% and 51% in CF-CF and CF-SS, respectively (Fig. S8). At the genus level, the abundance of *Methanobacterium* was almost similar (13-14%) in the anode biofilms in both systems (Fig. 5b). However, the abundance of *Methanobacterium* in cathode biofilms of CF-SS was higher than CF-CF (51% vs. 32%). Previous studies also reported the enrichment of known hydrogenotrophic methanogens in methanogenic biocathode ^8–11^. Moreover, *mcr*A gene sequencing was performed (Fig. 5c) to understand the taxonomy of methanogens ^53,70,71^. In the anodic biofilms, the abundances of *Methanobacterium* species were almost similar, including *formicicum* (CF-CF: 67%; CF-SS: 71%) and *subterraneum* (CF-CF: 34%; CF-SS: 29%). In the cathodic biofilms, CF-SS showed more diverse species of *Methanobacterium*; *formicicum* (54%), *subterraneum* (24.4%), and *palustre* (22%), as compared to CF-CF; *formicicum* (81%), and *subterraneum* (19%). Thus, the higher abundance and diverse species of hydrogenotrophic *Methanobacterium* on stainless steel cathode might have contributed to the faster methanogenesis via hydrogen utilization.

#### 3.6.3. Principal component analysis

The PCA analysis of biocathode bacterial and archaeal communities was performed to evaluate the relation between genera and PCs (Fig. S9). Based on 16S rRNA bacterial sequencing of biocathode, the superior performance of CF-SS was related to the enrichment of homoacetogenic *Acetobacterium* (Fig. S9a). However, the other genera might have an indirect relation to the superior performance of CF-SS. Based on archaeal sequencing of biocathode, hydrogen-consuming *Methanobacterium* and *Acetobacterium* primarily contributed to the superior performance of stainless steel biocathode (Fig. S9b).

### 3.7. Expression of EET genes

The gene expression for *pil*A and c-type cytochromes (Fig. 6) shows trivial differences in their expression levels in anode/cathode biofilms between both reactors. Moreover, compared to anode biofilms, the EET-associated genes were less expressed in cathode biofilms in both reactors. Based on the authors’ knowledge, this study first reports the expression of EET genes for methanogenic biocathode. The EET from EAB to the anode has been demonstrated to be facilitated via c-type cytochromes and conductive nanowire or pili ^74^, while the significance of EET in methanogenic biocathode is still ambiguous. However, previous reports postulated that conductive pili and c-type cytochromes could play an important role in DIET from EAB to methanogens ^75,76^. Notably, some bacteria (e.g., *Enterobacteriaceae, Desulfovibrio*, etc.) found in biocathode in this study could express different cytochromes and/or conductive pili ^74,77^. Furthermore, a recent study suggested that *Methanobacterium* species could produce methane via DIET ^78^. Nonetheless, the expressions of these EET genes were quite comparable in both systems, indicating higher current density and methane productivity from the CF-SS reactor was not attributed to the overexpression of EET genes.

**Fig. 6.**
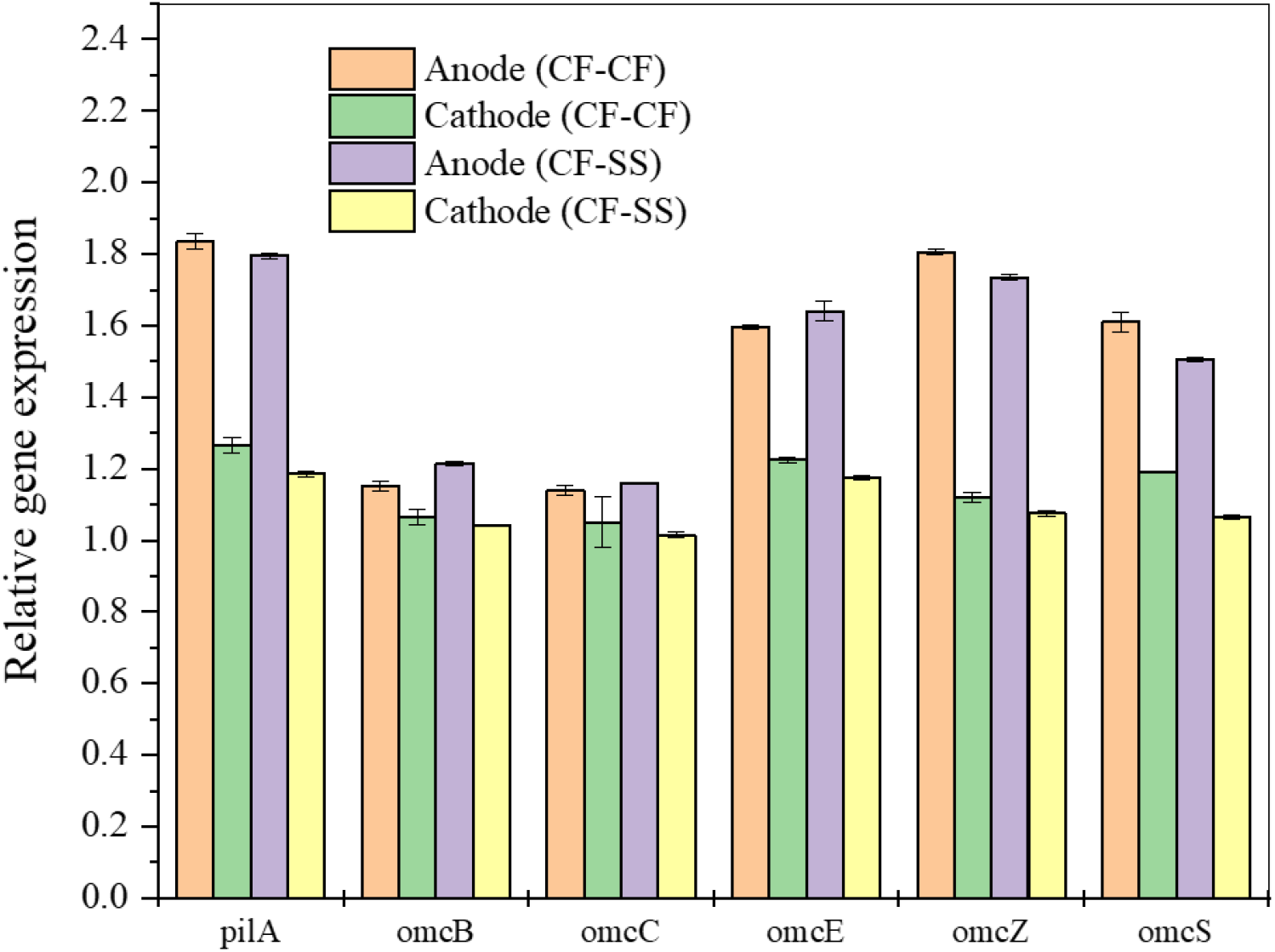
Expression of genes known to regulate extracellular electron transfer in biofilms. The error bars indicate the standard deviation of three replicates (n = 3).

### 3.8. Implications

This study provides new insights into the characteristics and significance of EPS and expressions of EET genes in methanogenic biocathode. As compared to the carbon fiber, significantly higher EPS levels were observed in the stainless steel biocathode. Protons reduction to H_2_ during HER can create local alkaline pH on the cathode. Thus, it could be posited that the highest EPS secretion in stainless steel biocathode could be linked with faster HER. One important finding of this current study is that EET may not play a decisive role in differentiating performances in MEC-AD systems using different electrode materials. Instead, the effective catalysis of HER, lower internal resistance, and higher abundances of H_2_-utilizing methanogens and homoacetogens on stainless steel cathode appeared to be the primary reason behind the higher methanogenic activity. Nonetheless, based on EET gene expression patterns and redox activity of biocathode-derived EPS, EET would still be involved in cathodic electro-methanogenesis.

Regarding the engineering significance of the results, carbon-based cathode electrodes have been mostly used in MEC-AD systems due to their excellent biocompatibility and higher surface area over metal-based electrodes ^3,30–32^. While carbon fibers provided a higher specific surface area, stainless steel mesh outperformed carbon fibers under similar operating conditions (e.g., anode electrode, inoculum, mixing, etc.). Given that most of the single-chamber MEC-AD studies used carbon-based biocathode ^5,7^, the results of this study are significant for selecting efficient cathode materials to realize improved performance. However, it should be noted that the results presented here are from specific operating conditions with two selected electrode materials. Hence, further research is warranted with more carbon and metal electrodes with similar textures and surface areas.

## Associated Content

### Supporting Information

Estimation of specific surface area of electrodes (Fig. S1); EPS extraction and analytical methods (Table S1); Method for CLSM imaging; CLSM imaging of ROS (Fig. S2); Methods for EET gene expression measurement (Table S2); Microbial quantification using RT-PCR; Current density (Fig. S3); Electrochemical impedance spectroscopy (EIS) (Fig. S4); Profiles of COD and VFAs (Fig. S5); SEM images of biofilms (Fig. S6); CV of EPS (Fig. S7); Microbial diversity and richness (Table S3); Microbial communities at the phylum level (Fig. S8); PCA results (Fig. S9).

## Acknowledgements

This work was supported by the Discovery Grant from Natural Sciences and Engineering Research Council of Canada (NSERC) and the Early Career Research Award from Future Energy Systems (FES). Basem Zakaria acknowledges Killam Trusts for Izaak Walton Killam Memorial Scholarship for his doctoral studies.

## Notes

The authors declare no competing financial interest.

