## Supplementary Information for "Characterization and significance of extracellular polymeric substances, reactive oxygen species, and extracellular electron transfer in methanogenic biocathode" (from another version of this preprint: version 1, 2020-11-14)

### New insights into biocathode material as a key determinant of electro-methanogenic activity and significance of extracellular polymeric substances

Basem S. Zakaria, Bipro Ranjan Dhar*

Civil and Environmental Engineering, University of Alberta, 9211-116 Street NW, Edmonton, AB, Canada T6G 1H9

**Text S1. Estimation of specific surface area of carbon fiber and stainless-steel mesh electrode**

**1. Stainless-steel mesh electrode**

The surface area was calculated according to Zhang et al. (2010) with minor modification.

$$S = 2\pi b d n (n+1) + \frac{3}{2}\pi \ d^2 (n+1)^2$$

Where, (b) is the pore size, and (d) is the wire diameter, (n) is 70 mesh

b= 0.02 cm, d = 0.016 cm

The specific surface area based on reactor volume = 0.0016 $m^2$/0.00038 $m^3$ = 4.23 $m^2/m^3$

**2. Carbon fiber electrode**

The surface area of carbon fiber was calculated using two approaches previously reported in the literature (Brunschweiger et al., 2020; Lanas and Logan, 2013):

1. Considering every single filament in the carbon fiber bundle (filament bundle consists of 24000 single filaments, each with diameter 7 μm, and the length = 130 cm)

S.A. = 2πrh x n = 2 x 3.14 x 0.0007 cm x 130 cm x 24000 = 13715.5 $cm^2$

The specific surface area based on reactor volume = 1.372 $m^2$ / 0.00038 $m^3$ = 3609 $m^2/m^3$

2. Considering all filaments in a bundle act together as a single fiber (width = 0.5 cm, length = 130 cm, height = 0.1 cm).

S.A.= 2 (wh +Lw + Lh) = 156.4 $cm^2$

The specific surface area based on reactor volume = 0.0156 $m^2$ / 0.00038 $m^3$ = 41 $m^2/m^3$

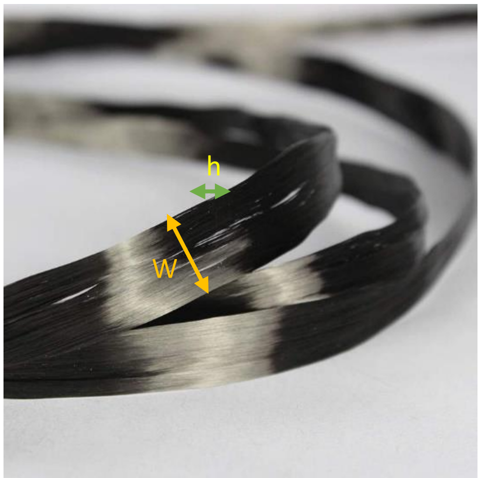


**Fig. S1.** Photograph of carbon fiber bundle.

**Text S2. EPS extraction protocol:** CER method for EPS extraction was performed according to Frølund et al. (1996) with some minor modification, 52 $cm^2$ of carbon fiber and 0.054 $cm^2$ of stainless-steel were resuspended in 0.1 M PBS buffer, pH 7.4. Then, 2 g CER (Dowex® Marathon® C sodium form, Sigma-Aldrich, USA), washed with 0.1 M PBS for 15 min (10 mL $g^{-1}$ Dowex), was added to each sample. The samples were agitated at a high capacity for 20 min on the shaker (Vortex Mixer, Fisher Scientific, USA). The heating method was conducted according to Xu et al. (2013), the electrodes were washed twice with 0.9% NaCl (w/v), then resuspended in 0.9% NaCl (w/v). The samples were heated at 60$^o$C in a water bath for 30 minutes. For both methods, the EPS were harvested by centrifugation three times at 20,000 × g for 20 minutes at 4$^o$C. The supernatant was collected and filtered using a 0.2 µm filter and the EPS stored at -20$^o$C for further analysis. Also, the pellets were collected to examine the cell lysis using Glucose-6-Phosphate Dehydrogenase kit (Sigma-Aldrich, USA).

**Table S2.** EPS composition using CER and heating methods.

| | Carbohydrate | Proteins | Heme | Uronic Acid | eDNA |
|---|---|---|---|---|---|
| CER Method | | | | | |
| **Anode (CF-CF)** | 28.225 | 133.545 | 40.58 | 10.6 | 0.55 |
| **Cathode (CF-CF)** | 25.775 | 170.025 | 16.52 | 6.2 | 0.39 |
| **Anode (CF-SS)** | 34.775 | 146.405 | 37.94 | 11.665 | 0.685 |
| **Cathode (CF-SS)** | 52.14 | 212.79 | 34.22 | 15 | 1.525 |
| Heating Method | | | | | |
| **Anode (CF-CF)** | 35.575 | 109.29 | 50.45 | 13.065 | 0.69 |
| **Cathode (CF-CF)** | 27.78 | 163.29 | 18.62 | 8.27 | 0.365 |
| **Anode (CF-SS)** | 24.1 | 172.855 | 41.13 | 11.32 | 0.46 |
| **Cathode (CF-SS)** | 52.61 | 223.885 | 37.57 | 15.2 | 1.715 |

**Text S3. Analytical methods for different EPS components:** The total carbohydrate content was measured using a phenol-sulfuric acid method using glucose standards according to DuBois et al. (1956) with some minor modification. 2 mL of EPS was mixed with 5 mL of concentrated sulfuric acid followed by the addition of 0.05 mL of 80% wt. phenol. Then, the samples were incubated at room temperature for 10 minutes, then shake followed by incubation for 20 minutes at 30$^{o}$C. The samples were measured at 490 nm using UV-spectrophotometer (Model DR 3900, HACH, Germany). The protein contents of the EPS were measured using Pierce™ Modified Lowry Protein Assay Kit (Thermo Fisher, USA) according to the manufacturer's instructions. Uronic acid was measured according to Blumenkrantz and Asboe-Hansen (1973) and Ghods et al. (2015) with glucuronic acid (Sigma-Aldrich, USA) as a standard. Heme contents were measured using the Heme Assay Kit (Heme Assay Kit, Sigma-Aldrich, USA) according to the manufacturer's instructions. The extracellular DNA (eDNA) was quantified using the diphenylamine colorimetric method with calf thymus DNA as the standard (Brunk et al., 1979). 100 µL EPS was pipetted in a cuvette and 2.5 mL of the reagent (0.2 ppm DAPI (4, 6-diaminodino-2-phenylindole) in 100 mM NaC1, l0 mM EDTA, 10 mM PBS, pH 7.2) was added. All the above measurements were conducted in triplicate.

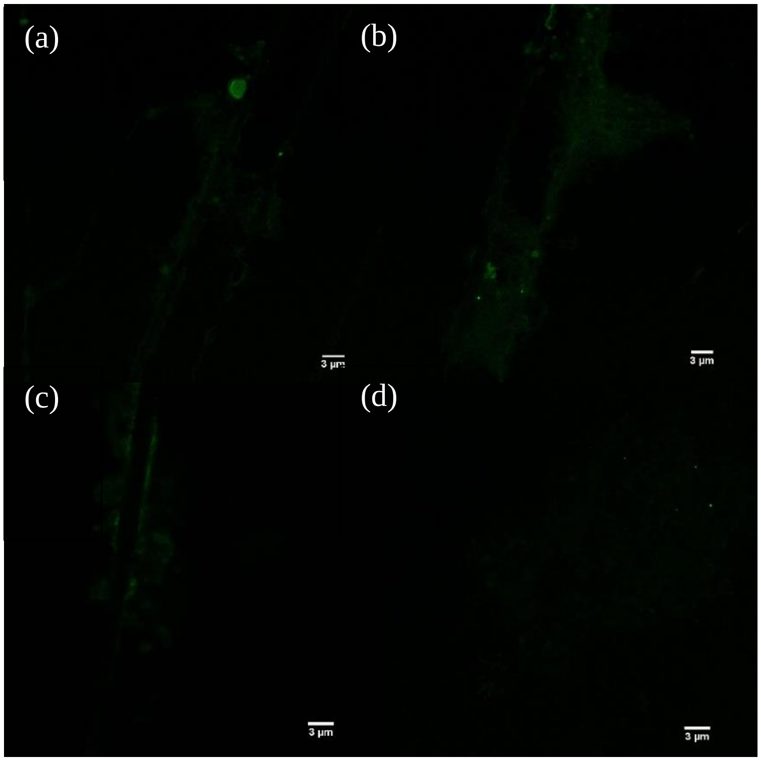


**Fig. S2.** CLSM of reactive oxygen species (ROS). Anode (CF-CF) (a), cathode (CF-CF) (b), anode (CF-SS) (c), and cathode (CF-SS) (d).

**Text S4. RT-PCR of gene expression levels.** The concept is to design primers to detect most of the *omc*B, *omc*C, *omc*E, *omc*Z, *omc*S, and *pil*A gene sequences as possible. The primer design was performed similarly to Lin et al. (2019), and *rec*A housekeeping gene was used as a reference (Rivas et al., 2005). The genome sequences of the anode biofilm were collected from the National Center for Biotechnology Information (NCBI) (https://www.ncbi.nlm.nih.gov). These followed by multiple-aligned using the ClustalX alignment tool (http://www.clustal.org/clustal2/) to select most of the homologous gene regions (Thompson, 1997). Then, gene-specific primers for RT-PCR were designed with Primer3 software (http://bioinfo.ut.ee/primer3) by selecting an almost similar range of melting temperatures. The Basic Local Alignment Search Tool (BLAST) (https://blast.ncbi.nlm.nih.gov/Blast.cgi) was used to check the specificity of the primers. Also, the primers were experimentally examined using agarose gel electrophoresis. The primers were prepared by Integrated DNA Technologies (IDT, USA) and are listed in Table S1.

**Table S1.** Primers used for studying gene expression levels

| | Forward (5'-3') | Reverse (5'-3') |
|---|---|---|
| *rec*A | CACCGGCATAATCTCCAAGT | ATCTTGCGGATATCGAGACG |
| *pil*A | TTATGATCGTGGTTGCCATT | CCGCAGTTAAACCTTCTGCT |
| | GAGCGATGTTCTTTCCGTTT | AATCCTGATAGGCGGGAATC |
| omcB | CCGAAAATTACGCAGGTGTT | GGAGTTCACGAAACCAAGGA |
| | GAGGGACGATGTCAACCTGT | TTCGCAAGGTAGCTGTTGTG |
| omcC | CTGGTACTCTGGGTGGCATT | GCTGTGGTTAGCAGCATTGA |
| | ACGAGTTCCAGACCAACACC | GGCAGTATCGTCCCAGTTGT |
| omcE | ATGTCCTACGGCGATGCTAT | GTAACCTGCAGGAAGGTGGA |
| | CTCGTCCAGCAGCATGAATA | GGGGTGATCATTGCTCAGAT |
| omcZ | GGACGTATTGTGGCAGAGGT | GGCCACTACATTCCGACCTA |
| | GGCCACTACATTCCGACCTA | GGACGTATTGTGGCAGAGGT |
| omcS | GCTGACTACACCGTGCTCAA | TGAACTCGTATGCCAGGTTG |
| | GGCAAGTACCGTCGTTTTGT | GTAGCTTCCGTCCGGTTGTA |

**Text S5. Microbial quantification using RT-PCR.** qPCR was performed to quantify bacterial cell numbers before and after AgNPs injection. 16S universal primers, 357Wf: CCTACGGGNGGCWGCAG and 785R: GACTACHVGGGTATCTAATCC, were used to quantify the DNA samples. qPCR mixtures were prepared in 25 uL reactions using QuantiFast SYBR® Green PCR Kit (Qiagen, CA) as the following: 2 µL of the DNA template, 12.5 µL 2x master mix, 2.5 µL forward and reverse specific primer, and 5.5 µL nuclease-free water. CFX 96 real-time PCR system with a C1000 Thermal Cycler (Bio-Rad, USA) was used with the following cycling conditions according to the QuantiFast SYBR® Green PCR Kit's protocol; PCR initial heat activation cycle at 95 ˚C for 5 min, 35 cycles at 95 ˚C for 10 sec and 60 ˚C for 30 sec, and finally, one cycle at 40 ˚C for 30 seconds. Triplicate reactions were run for all samples.

**Table S3.** The diversity and richness of the biofilms of both systems.

| Reactor/electrode | | Chao 1 | Phylogenetic distance | OTUs | Pielou's evenness | Shannon | Coverage |
|---|---|---|---|---|---|---|---|
| **CF-CF** | **Anode** | 137 | 11.68 | 132 | 0.45 | 3.14 | 1.0 |
| | **Cathode** | 76 | 7.19 | 76 | 0.63 | 3.95 | 1.0 |
| **CF-SS** | **Anode** | 146 | 13.57 | 146 | 0.72 | 5.18 | 1.0 |
| | **Cathode** | 155 | 12.32 | 150 | 0.71 | 5.10 | 1.0 |

(a)

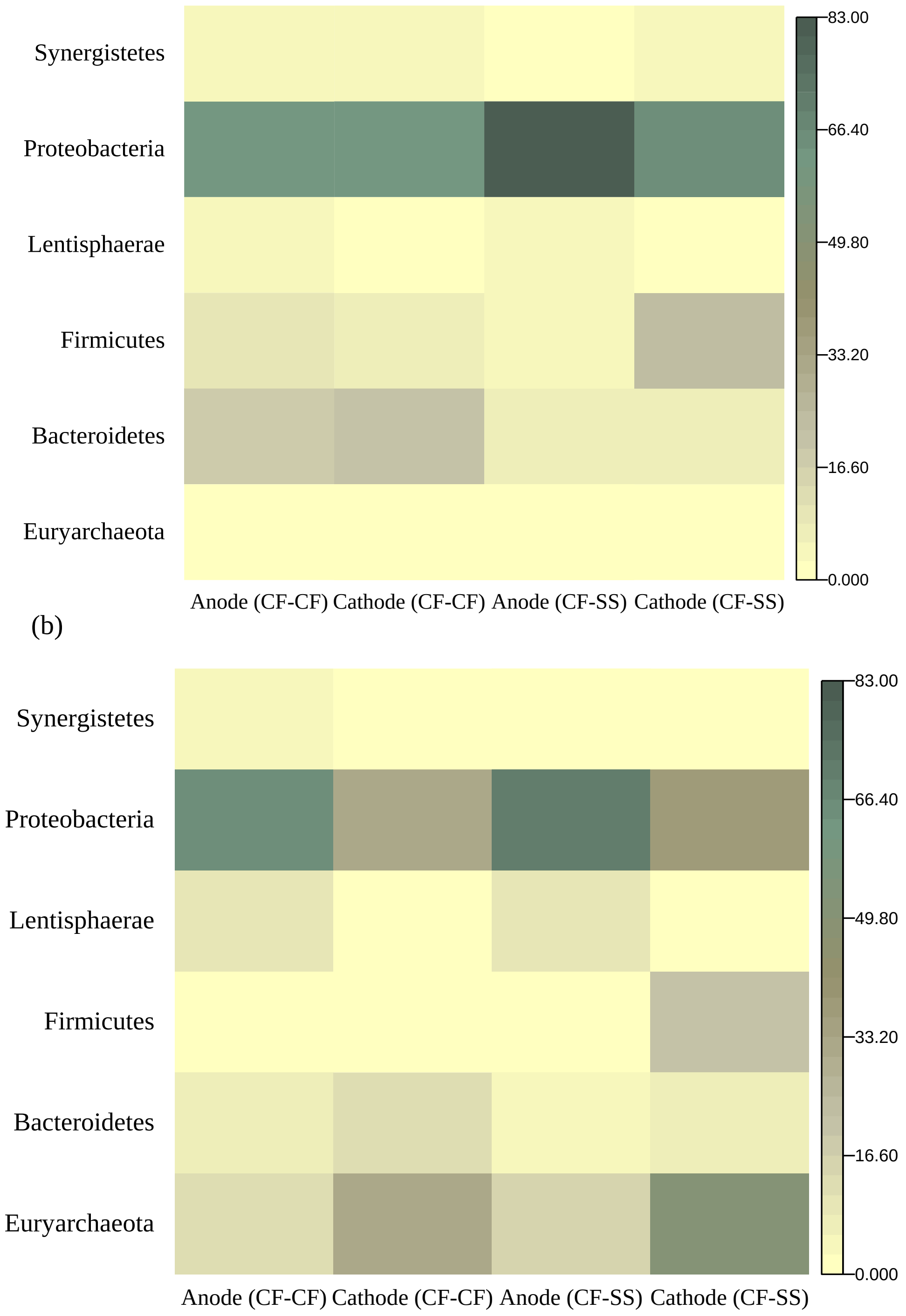


**Fig. S3.** Relative abundance of microbial community. Bacterial primer phylum level (a), and archaeal primer phylum level (b).
